# Asymmetric Contribution of a Selectivity Filter Gate in Triggering Inactivation of Ca_V_1.3 Channels

**DOI:** 10.1101/2023.09.21.558864

**Authors:** Pedro J. del Rivero Morfin, Audrey L. Kochiss, Klaus R. Liedl, Bernhard E. Flucher, Monica L.I. Fernández-Quintero, Manu Ben-Johny

## Abstract

Voltage-dependent and Ca^2+^-dependent inactivation (VDI and CDI, respectively) of Ca_V_ channels are two biologically consequential feedback mechanisms that fine-tune Ca^2+^ entry into neurons and cardiomyocytes. Although known to be initiated by distinct molecular events, how these processes obstruct conduction through the channel pore remains poorly defined. Here, focusing on ultra-highly conserved tryptophan residues in the inter-domain interfaces near the selectivity filter of Ca_V_1.3, we demonstrate a critical role for asymmetric conformational changes in mediating VDI and CDI. Specifically, mutagenesis of the domain III-IV interface, but not others, enhanced VDI. Molecular dynamics simulations demonstrate that mutations in distinct selectivity filter interfaces differentially impact conformational flexibility. Furthermore, mutations in distinct domains preferentially disrupt CDI mediated by the N- versus C-lobes of CaM, thus uncovering a scheme of structural bifurcation of CaM signaling. These findings highlight the fundamental importance of the asymmetric arrangement of the pseudo-tetrameric Ca_V_ pore domain for feedback inhibition.

## Introduction

The high voltage-activated Ca^2+^ channels (Ca_V_1/2) serve as vital conduits for Ca^2+^ entry responsible for orchestrating diverse physiological processes, ranging from excitation-contraction coupling in muscle (Bers, 2002), to vesicle secretion and gene transcription in neurons (Berridge et al., 2000; Dolmetsch, 2003). Molecularly, the eukaryotic Ca_V_1/2 channels possess a conspicuous pseudo-tetrameric architecture that is thought to permit asymmetric conformational changes to fine-tune channel activity (Catterall et al., 2017). For related Na_V_ channels, this asymmetric architecture allows distinct voltage-sensing domains (VSDs) to differentially impact activation versus inactivation and pore-conformation (Goldschen-Ohm et al., 2013). By comparison, although Ca_V_ channels have VSDs with distinct kinetics, the importance of the overall pseudo-tetrameric architecture and potential asymmetric conformational changes are not fully established, though likely important (Savalli et al., 2021). Like Na_V_, Ca_V_ channels are subject to exquisite feedback regulatory mechanisms, including both voltage-dependent inactivation (VDI) and Ca^2+^-dependent inactivation (CDI) that limits excessive Ca^2+^ entry into the cell (Minor and Findeisen, 2010; Ben-Johny and Yue, 2014). Both processes are biologically consequential, as Ca^2+^ channelopathies involve deficits in inactivation that disrupt electrical signaling and Ca^2+^ homeostasis, thereby contributing to neurological, neurodevelopmental, and cardiac arrhythmogenic disorders such as Timothy Syndrome (TS) (Barrett and Tsien, 2008; Dick et al., 2012; Limpitikul et al., 2014; Pinggera et al., 2015). Both inhibitory processes have disparate molecular origins. VDI is initiated by voltage-dependent conformational changes triggered by the movement of the four homologous transmembrane voltage-sensing domains (Stotz and Zamponi, 2001; Stotz et al., 2004; Tadross et al., 2010). By contrast, CDI is largely independent of voltage (Tadross and Yue, 2010) and, instead, ensues from Ca^2+^ binding to the two lobes of calmodulin (CaM) tethered to the Ca_V_ carboxy-terminus (Peterson et al., 1999; Qin et al., 1999; Zuhlke et al., 1999). These processes converge on distinct ‘gates’ in the pseudo-symmetric pore domain to obstruct ion influx (Tadross et al., 2010; Abderemane-Ali et al., 2019). Even so, the exact nature of conformational changes that underlie channel feedback inhibition, and the contribution of the pseudo-symmetric channel architecture are not fully understood.

Three end-stage mechanisms of inactivation have been proposed, paralleling findings with related ion channels (Fig. 1A). These include: (1) a hinged-lid mechanism that involves a cytosolic inactivation particle that occludes the ion conduction pathway (Stotz et al., 2000) similar to N-type inactivation of Shaker K^+^ channels (Hoshi et al., 1990) or fast inactivation of voltage-gated Na^+^ (Na_V_) channels (West et al., 1992; Yan et al., 2017), (2) an allosteric closure or destabilization of the S6 activation gate (Imredy and Yue, 1994; Tadross et al., 2010) similar to allosteric regulation of K_V_7 channels (Zaydman and Cui, 2014), and (3) a conformational rearrangement of the selectivity filter (SF) (Abderemane-Ali et al., 2019), akin to P-type (or C-type) inactivation in Shaker K^+^ channels (Hoshi et al., 1991) or slow inactivation of Na_V_ channels (Balser et al., 1996). For VDI, a hinged-lid mechanism has been proposed (Scheme 1), with the N-terminus of the intracellular domain I – II (DI-DII) linker serving as an inactivation particle that binds to a transmembrane receptor site formed by the S6 gate (Stotz et al., 2000; Dafi et al., 2004; Tadross et al., 2010). For CDI, initial studies indicated an allosteric mechanism, where channels switch to a distinct gating pattern with sparse openings (Scheme 2) (Imredy and Yue, 1994; Tadross et al., 2010). More recently, structure-guided mutagenesis experiments point to the involvement of an alternate SF gate for both VDI and CDI (Scheme 3) (Abderemane-Ali et al., 2019). Even so, the extent of SF conformational changes in channel feedback inhibition and key factors that govern this process remains to be fully understood.

**Figure 1.**
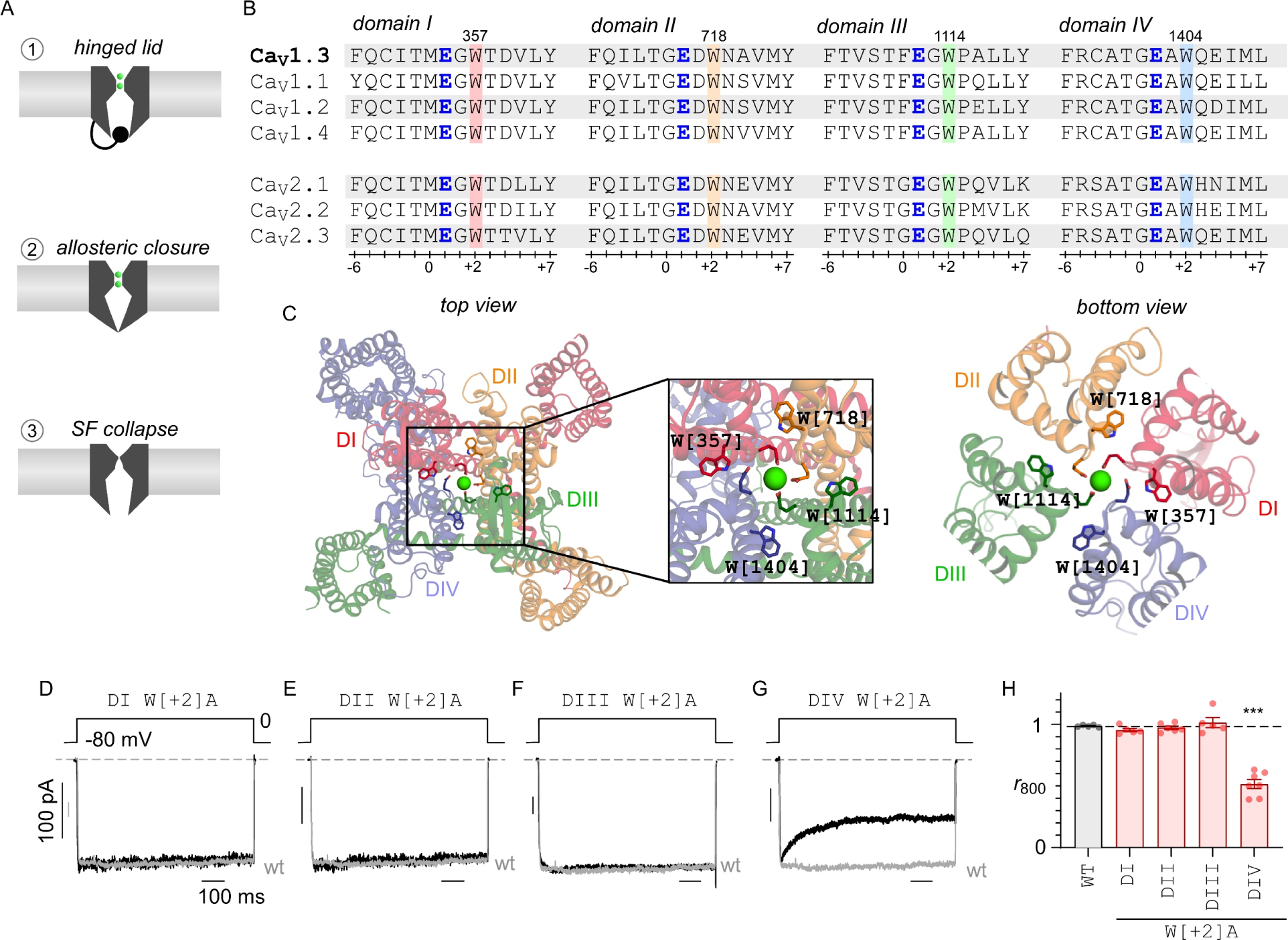
Asymmetric role of Domain IV pore helix in tuning VDI of Ca1.3. (**A**) Three distinct end-stage mechanisms for Ca_V_1.3 inactivation. Scheme 1 shows a hinged-lid mechanism where the pore is occluded by an inactivation particle (DI-DII linker), Scheme 2 shows an allosteric mechanism where the open probability is reduced through increased channel closure, and Scheme 3 shows selectivity filter collapse. **(B)** Sequence alignment shows the conservation of a tryptophan residue (W) in the pore segment adjacent to the putative Ca^2+^ coordinating glutamate (E) residue. **(C)** Structural models show the pseudo-tetrameric architecture of Ca_V_1.3 (7UHG). Inset shows the ion conduction pathway lined by the selectivity filter Ca^2+^ coordinating glutamate residues and the spatial location of the conserved tryptophan. **(D)** Representative Ba^2+^ current traces evoked in response to an 800 ms voltage-step to 0 mV show minimal VDI of both wild-type (gray) and DI W[+2]A mutant channels. The wild-type current is normalized to compare inactivation kinetics. **(E-F)** Similarly, both DII W[+2]A and DIII W[+2]A mutant channels also showed minimal VDI. Format as in Panel C. **(G)** By contrast, DIV W[+2]A mutation enhanced VDI of Ca_V_1.3. Format as in Panel C. **(H)** Population data shows fraction of remaining current following 800 ms depolarization (*r*_800_). The DIV W[+2]A mutation selectively increases inactivation in comparison to both wild-type (gray) and W[+2]A substitutions in DI-DIII. Each bar, mean ± s.e.m. Wild-type, *n* = 5 cells, DI W[+2]A, *n* = 5 cells, DII W[+2]A, *n* = 6 cells, DIII W[+2]A, *n* = 5 cells, DIV W[+2]A, *n* = 7 cells. \*\*\**p* < 0.001 by Tukey’s multiple comparisons test.

Here, we sought to delineate the functional importance of conformational changes in the SF in mediating both VDI and CDI. We focused on Ca_V_1.3 channels for two reasons. First, these channels exhibit characteristically slow VDI kinetics owing to the presence of a “shield” that counteracts a hinged-lid inactivation mechanism (Tadross et al., 2010). This permits a bottom-up approach, whereby VDI may be engineered, thereby allowing us to probe the functional role of the SF gate. Second, these channels also demonstrate robust CDI that depends on distinct conformational changes involving both the N- and C-lobes of CaM. We considered an ultra-highly conserved Trp residue (W[+2]) within the SF found at the interface between the pore helices from different domains (Payandeh and Minor, 2015) (Fig. 1B-C). Systematic mutagenesis of the corresponding residue in each domain revealed an asymmetric switch in the pore domain that unveils VDI. Furthermore, in depth analysis of CDI showed that mutating the W[+2] residue in distinct domains asymmetrically diminished CDI mediated by distinct CaM lobes. In all, these findings highlight the fundamental importance of the asymmetric arrangement of the pseudo-tetrameric Ca_V_1.3 pore domain for channel inactivation.

## Results

### Mutations in the pore domain tune inactivation of Ca_V_1.3

Previous studies have identified the presence of a highly conserved Trp (W[+2]) residue two residues downstream of the selectivity filter of Ca_V_, Na_V_, and NALCN channels (Payandeh and Minor, 2015). Fig. 1B shows sequence alignment confirming the preservation of this residue within all four domains of Ca_V_ channels, while Fig. 1C shows its structural location within the inter-domain interfaces. We individually substituted W[+2] residues in DI–DIV of Ca_V_1.3 with alanine, yielding W[357]A, W[718]A, W[1114]A, and W[1404]A mutant channels respectively. For convenience, we refer to these mutations by the nomenclature: D*x* W[*y*]A where *x* denotes the particular domain and *y* refers to the position relative to the central Glu that forms the SF. To probe changes in voltage-dependent inactivation (VDI), we transfected either wild-type or mutant channels in HEK293 cells and undertook whole-cell current recordings utilizing Ba^2+^ as the charge carrier. Here, we co-transfected both α_2_δ and β_2A_ subunits, as these subunits are essential for trafficking and proper function of the Ca_V_1.3 channel. Importantly, the β subunit is well known to tune inactivation properties. As such for these experiments, we chose β_2A_, a palmitoylated variant that strongly reduces inactivation compared to other β subunit isoforms (Olcese et al., 1994; Chien et al., 1996; Colecraft et al., 2002; Van Petegem et al., 2004; He et al., 2007). As Ba^2+^ binds poorly to calmodulin (Chao et al., 1984), this maneuver permits quantification of VDI independent of Ca^2+^/CaM-dependent inactivation (CDI). For the wild-type Ca_V_1.3 short variant, an 800ms voltage-step to 0 mV revealed rapid activation and minimal inactivation, consistent with previous studies (Fig. 1D, gray trace). Similarly, analysis of DI W[+2]A, DII W[+2]A, and DIII W[+2]A mutant channels also revealed minimal VDI (Fig. 1D-F, black traces). By contrast, exemplar current recordings with the DIV W[+2]A mutant show a rapid decay of Ba^2+^ currents following initial activation, demonstrating increased inactivation (Fig. 1G, black trace). To quantify the steady-state extent of inactivation, we computed the ratio of the current remaining following 800ms of depolarization with that of the peak value (*r*_800_). Population data confirmed a significant increase in inactivation with the DIV W[+2]A mutation, manifesting as a reduced *r*_800_ value, in comparison to wild-type and other mutant channels (Fig. 1H). For all mutants, we observed minimal changes in the normalized current-voltage relationship suggesting minimal effect of the SF mutations on channel activation (Fig. S1). These findings suggest that the pore domain constitutes a key structural determinant for VDI in Ca_V_1.3. Furthermore, the selective effect of DIV W[+2]A in enhancing VDI highlights the functional asymmetry of pore domain conformational changes.

### Distinct functional signature of SF-dependent VDI of Ca_V_1.3

Canonical VDI of Ca_V_1/2 channels follow a boltzmann relationship with voltage, whereby currents evoked at more depolarizing potentials exhibit stronger inactivation. We, therefore, sought to determine whether enhanced Ca_V_1.3 VDI resulting from the pore-domain mutation also follows a similar trend. As such, inactivation of Ca_V_1.3 DIV W[+2]A mutant channels was determined by measuring fractional decay of whole-cell Ba^2+^ currents evoked in response to a family of step-depolarizations (Fig. 2A-B). Normalizing each current trace to its peak value showed that the extent of inactivation was reduced for DIV W[+2]A mutant at higher potentials, but not wild-type channels (Fig. 2A-B, Fig. S2A). To further quantify the voltage-dependence of inactivation, we utilized a two-pulse protocol (Fig. 2C-F) (Patil et al., 1998). A brief 15 ms pre-pulse to 10 mV is used to probe the available current prior to VDI (*I*_pre_) (Fig. 2C, Fig. S2B). A family of 800 ms step-depolarizations to various test-pulse potentials (*V*_test_) is then used to elicit steady-state levels of inactivation, while a 15 ms post-pulse to 10 mV measures the residual current, reflecting ion influx through non-inactivated channels. The strength of VDI is quantified as the ratio of peak current during the post-pulse (*I*_post_) to that during the pre-pulse (*I*_pre_) (Fig. S2B). For wild-type Ca_V_1.3, we observed a modest increase in peak currents during the post-pulse as compared to the pre-pulse at more depolarized test-potential, indicating the presence of weak voltage-dependent facilitation (Fig. 3C,E; Fig. S2B). By comparison, for the DIV W[+2]A mutant, we observed a U-shaped voltage-dependence of VDI, with the largest reduction in peak current observed with modest depolarizations (Fig. 3D,F; Fig. S2C). We further assessed voltage-dependence of inactivation with either Ca^2+^ or Na^+^ as charge carrier. In both cases, we observed a U-shaped voltage-dependence (Ca^2+^, Fig. S2D-F; Na^+^, Fig. S2G-I). Of note, for these experiments, we co-expressed Ca^2+^-insensitive mutant CaM (CaM_1234_) whose Ca^2+^ binding sites in its EF hands were disabled (Xia et al., 1998; Peterson et al., 1999). As maneuver disables CDI, it allow us to dissect CaM independent effects of Ca^2+^ on channel gating.

**Figure 2.**
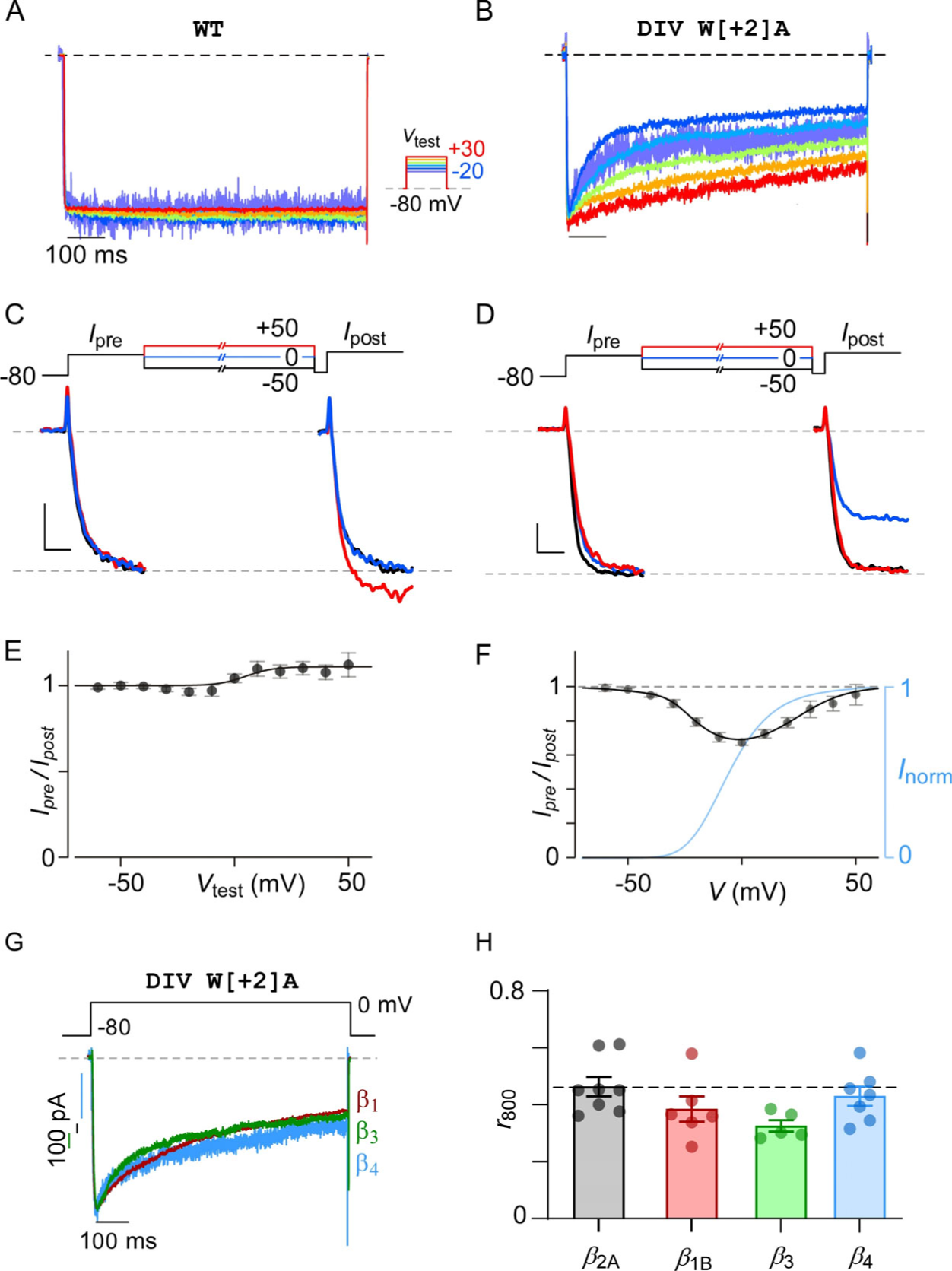
Distinct functional signature of Ca_V_1.3 VDI. **(A-B)** Exemplar Ba^2+^ current traces evoked in response to a family of voltage-steps show minimal VDI of both wild-type Ca_V_1.3 (panel A) and DIV W[+2]A mutant (panel B). CaM_1234_ co-expressed to exclude potential contribution of CDI. **(C)** A two-pulse protocol is used to quantify voltage-dependence of inactivation. Comparison of Ba^2+^ currents for wild-type Ca_V_1.3 during the pre-pulse versus the post-pulse. Ca_V_1.3 currents are increased in response to higher test-pulse potentials. See Fig. S2B for full exemplar trace. (**D**) DIV / W[+2]A mutant channels exhibit diminished peak currents during the post-pulse following test pulse to intermediate voltages. See Fig. S2C for full exemplar trace. (**E**) Analysis of wild-type Ca_V_1.3 reveals minimal VDI across multiple voltages. Each dot and error, mean ± s.e.m. *n* = 7. (**F**) VDI of DIV / W[+2]A mutation exhibits a U-shaped voltage-dependence with strongest inactivation observed near 0 mV. Blue curve, voltage-dependence of inactivation obtained from whole cell normalized IV relations (Fig. S2E). Each dot and error, mean ± s.e.m. *n* = 7 cells. (**G**) Exemplar traces show similar kinetics of inactivation for DIV / W[+2]A mutant in the presence of various β subunits. **(H)** Population data confirms minor differences in VDI quantified as *r*_800_ measured at 0 mV in the presence of various β subunits. Each bar and error, mean ± s.e.m. *n* = 6 cells (β_1B_), *n* = 8 cells (β_2A_), *n* = 5 cells (β_3_), *n* = 7 cells (β_4_). *p* = 0.076 by one-way ANOVA.

**Figure 3.**
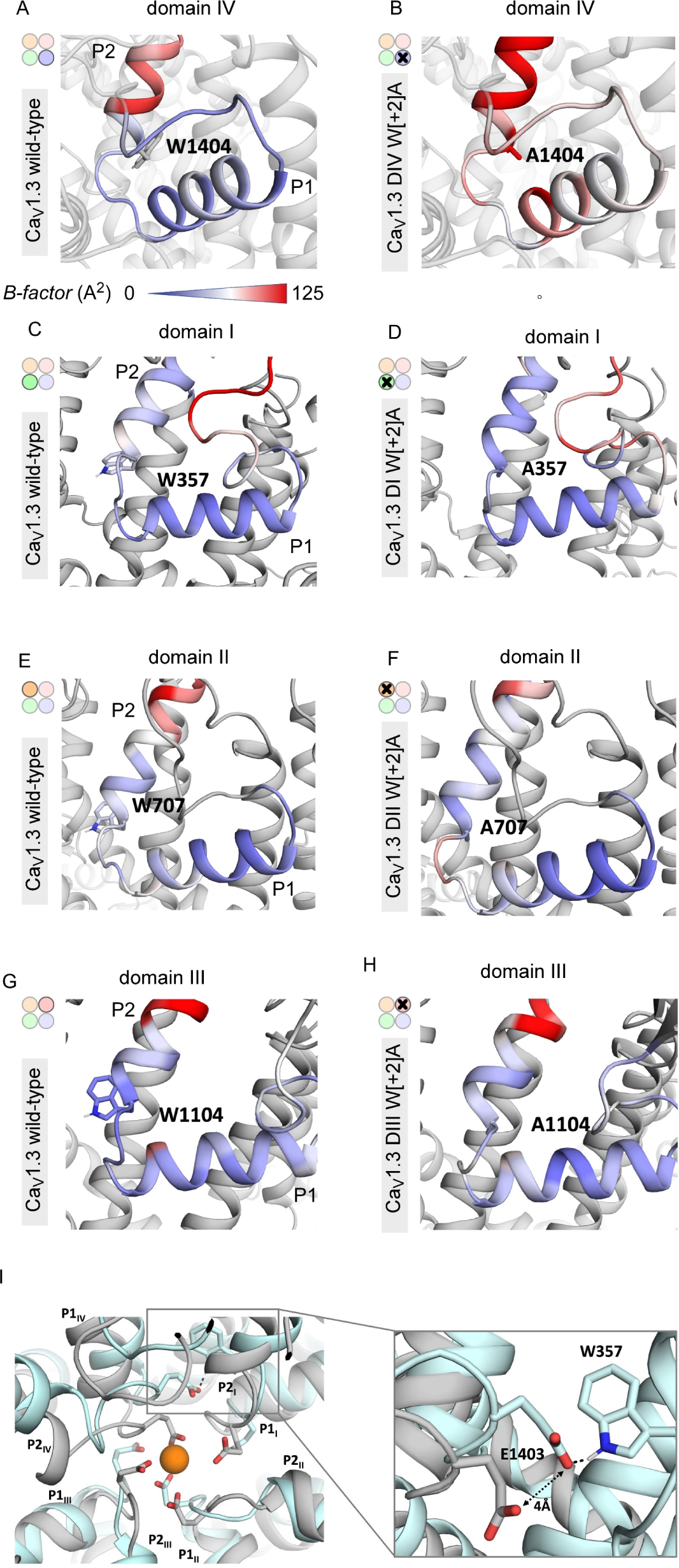
DIV W[+2]A mutation increases local conformational flexibility near the selectivity filter. **(A)** MD simulations were used to estimate local conformational flexibility of the domain IV pore helices of wild-type Ca_V_1.3 embedded in a lipid bilayer. Colormap indicates computed B-factors for residues in DIV pore helices ranging from 0 (green) to 147.7 (red) corresponding to increasing flexibility. **(B)** DIV W[+2]A mutation increases local conformational flexibility. **(C-D)** In comparison to wild-type channels (panel C), **s**imulations of DI W[+2]A mutant (panel D) show minimal change in computed B-factors for residues in DII pore helices. (**E-F**) Both wild-type (panel E) and DII W[+2]A mutant (panel F) show minimal change in local conformational flexibility. (**G-H**) Similarly, both wild-type (panel G) and DII W[+2]A mutant (panel H) exhibit minimal change in local conformational flexibility. This differential effect between DI-III versus DIV points to a specific role for DIV in tuning VDI. (**I**) Left, comparison of wild-type versus DIV W[+2]A mutation reveal key changes in the selectivity filter conformation. The DIV W[+2]A mutation alters the arrangement of Ca^2+^ coordinating E[0] residue, with DIV E[0] residue shifted by ∼ 4 Å. Right, detailed view of the DIV E[0] residue (E1402).

Conventional VDI of Ca_V_1/2 channels is highly tuned by the auxiliary β-subunit associated with the channel. More specifically, membrane-localized β-subunits (e.g., β_2A_) reduce VDI, while other isoforms yield strong VDI. For Ca_V_1.3, we previously identified mutations in the distal S6 gate that enhanced VDI (Tadross et al., 2010). However, the magnitude of VDI for these mutations varied depending on the specific β-subunit co-expressed. Here, to determine whether VDI conferred by SF mutation is also tuned by the auxiliary β-subunit, we co-expressed Ca_V_1.3 DIV W[+2]A mutant with non-membrane localized *β*_1_, *β*_3_, or *β*_4_ subunits. In all cases, we found that the extent of VDI was not further accentuated compared to that with the *β*_2A_ subunit (Fig. 2G-H). Taken together, these findings suggest that VDI conferred by SF mutation is a biophysically distinct process compared to conventional VDI of Ca_V_1 and Ca_V_2 channels.

### DIV W[+2]A mutation increases conformational flexibility of the SF

Given the asymmetric effect of DIV W[+2] residue and its location at the interface of two domains, we considered whether this mutation destabilizes the conformation of the SF. To probe this possibility, we performed molecular dynamics (MD) simulations (3x300 ns) based on the recently published Cryo-EM structure of the Cav1.3 channel of the WT and the mutants, embedded in a plasma membrane. To quantify potential changes in flexibility upon substitution of Tryptophan residues with alanine, we used the obtained simulations to calculate residue-wise B-factors. B-factors are an alignment-dependent measure to identify areas with higher fluctuations/flexibility. In wild-type channels the two helices of the pore loop are all very rigid (low B-factors; Fig. 3, A,C,E,G, green), except in domain IV, where the P1 helix displays elevated flexibility (high B-factor, Fig. 3A, red). In comparison to the wild-type channels, introduction of the DIV W[+2]A mutation increased the local conformational flexibility of this region evident as increased B-factor mapped on the structural model, particularly in the P2 helix (Fig. 3A-B). This suggests that this mutation destabilizes the SF conformation to potentially initiate inactivation. To evaluate this possibility further, we considered whether like mutations in the other domains also yielded increased conformational flexibility. In contrast to the DIV W[+2]A, we found that W[+2]A mutations in all other repeats did not appreciably increase the structural variability of the SF and only minimally altered the B-factors (Fig. 3C-H) suggesting that the conformational stability of the SF is largely preserved by these mutations. Apparently, the increased conformational flexibility of the SF of domain IV is not the direct consequence of the amino acid substitution in the P2 helix, rather it depends on the specific environment of the SF in this domain. Interestingly, further scrutiny of the DIV W[+2]A mutant revealed key alterations in the conformation of the Ca^2+^ coordinating E[0] residues that line the SF (Fig. 3I). Specifically, the DIV E[0] residue shifts away from the center of the ion permeation pathway by ∼4Å forming a contact with DI W[+2] residue (W357). This change likely prevents the high affinity coordination of a Ca^2+^ ion in the SF, ultimately impacting ion influx through the channel. Taken together, these results highlight the asymmetric effect of SF mutations on conformational stability and the unique role of DIV W[+2] mutation for VDI of Ca_V_1.3.

### Bulkiness of the selectivity filter DIV W[+2] residue tunes inactivation

Having inferred that the DIV W[+2]A mutant undergoes a distinct form of inactivation, we sought to dissect its underlying mechanisms. Structurally, the W[1404] residue is wedged at the interface between selectivity filter helices from domain IV and domain III (Fig. 1C). The alanine mutation would remove this wedge and potentially promote conformational changes in the selectivity filter that manifests as channel inactivation. As such, we reasoned that the bulkiness of the W[1404] side chain may be essential to stabilize the selectivity filter and tune inactivation kinetics. We thus replaced the tryptophan residue with a threonine (DIV W[+2]T) and undertook whole-cell recordings. We found that DIV W[+2]T also increased VDI; however, the net magnitude of increase was blunted in comparison to the DIV W[+2]A (Fig. 4A), consistent with the intermediate size of the threonine side-chain. In like manner, substitution of the DIV W[+2] residue with a valine (DIV W[+2]V) also yielded an intermediate increase in VDI (Fig. 4B). By contrast, replacing the DIV W[+2] residue with a bulky phenylalanine residue (DIV W[+2]F) yielded no appreciable increase in VDI as compared to wildtype channels (Fig. 4C). These findings qualitatively confirm the trend that increased bulkiness of the DIV W[+2] residue reduces the magnitude of VDI. To quantify this trend, we plotted the *r*_800_ values for various DIV W[+2] substitutions as a function of the side-chain accessible surface area (ASA). We observed a Boltzmann relationship between *r*_800_ and ASA, with substitution of bulkier residues resulting in diminished VDI (Fig. 4G). This overall relationship is consistent with the DIV W[+2] residue serving to stabilize the DIII –DIV SF interface.

**Figure 4.**
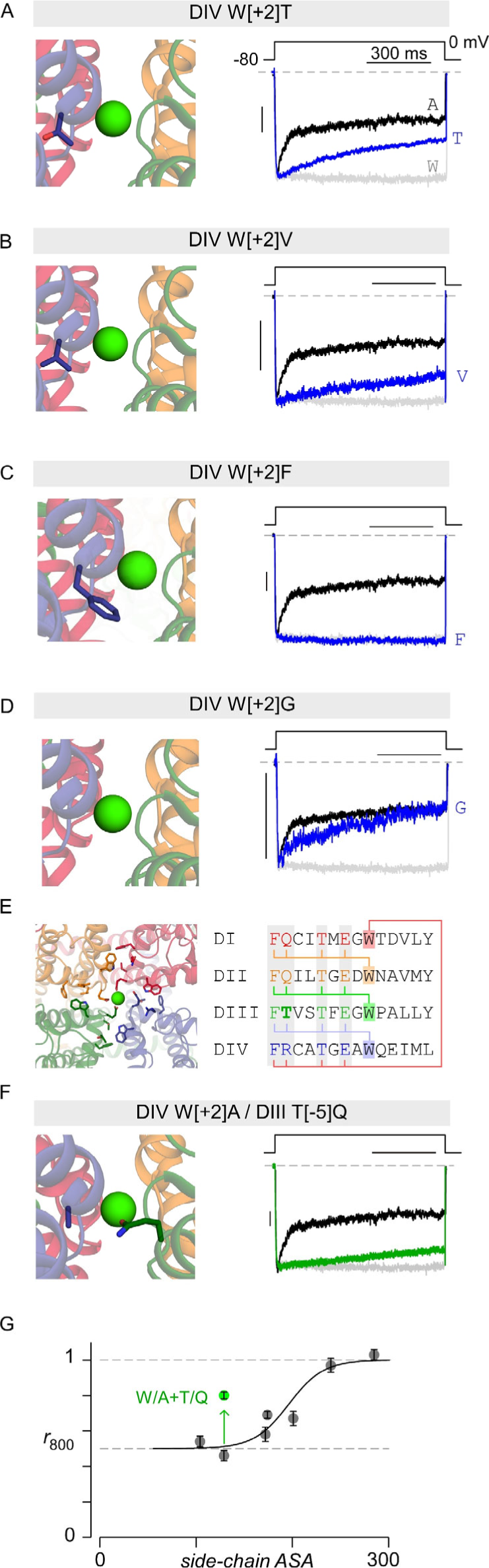
Bulkiness of the DIV W[+2] residue tunes VDI. **(A)** Exemplar traces reveal intermediate inactivation of Ca_V_1.3 DIV W[+2]T mutant (blue). Exemplar trances from wild-type (W[+2]) and DIV W[+2]A mutant channels are reproduced from Figure 1F to facilitate comparison. **(B)** VDI is similarly diminished for DIV W[+2]V mutation. Format as in panel A. **(C)** VDI is virtually absent for DIV W[+2]F mutation. (**D**) VDI is enhanced for DIV W[+2]G mutation. **(E)** Left, structural model highlights residues in adjacent domains that are within 5Å of W[+2] residue in a given domain. Right, sequence alignment and residues in close vicinity of W[+2] residue. Neighborhood residues E[0], T[-2], and F[-6] are conserved across different domains. By contrast, DIV W[+2] is in the vicinity of DIII T[-5] while either a Q or an R are present at the −5 position of other domains. **(F)** Analysis of DIII T[-5]Q / DIV W[+2]A double mutation shows reduced VDI in comparison to DIV W[+2]A single mutation. **(G)** Population *r*_800_ values for DIV W[+2] mutations reveal a Boltzmann relation between VDI as the accessible side-chain surface area. DIII T[-5]Q / DIV W[+2]A reduced VDI in comparison to DIV W[+2]A mutation and deviates from this relationship. Each symbol and errror, mean ± s.e.m from *n* = 5 cells for each mutant.

To further test this possibility, we hypothesized that introducing bulky side chains to residues in the DIII interface could compensate for the decreased stability resulting from DIV W[+2]A mutation. Accordingly, we identified candidate residues in domain III selectivity filter helices that are within 5 Å of the DIV W[+2] residue. These included residues F[1106] (DIII F[-6]), T[1107] (DIII T[-5]), T[1110] (DIII T[-2]), and E[1112] (DIII E[0]) (Fig. 4D). Examination of corresponding residues in domain I, II, and IV showed that F[-6], T[-2], E[0] are conserved. By contrast, the analogous position of DIII T[-5] residue in domains I, II, and IV is occupied by either a glutamine or an arginine (Fig. 4E). As such, we replaced T[1107] with a glutamine (T[1107]Q) in DIV W[+2]A mutant channel and probed changes in VDI. Indeed, we found that the DIV W[+2]A/DIII T[-5]Q double mutant exhibited substantially diminished inactivation as compared to DIV W[+2]A mutant channels (Fig. 4F-G). Taken together, these results suggest that the symmetric W[+2] residues determine the stability of the selectivity filter in concert with the asymmetric inter-domain interface residues. The two together are critical to prevent the collapse of the ion conduction pathway, and thus confer a unique role for DIV in suppressing VDI in Ca_V_1.3 channels.

### Pore domain mutations disrupt CDI in a lobe-specific manner

Distinct from VDI, Ca_V_1.3 channels undergo rapid Ca^2+^-dependent inactivation (CDI) mediated by calmodulin (CaM). Recent studies suggest that the end-stage mechanism of CDI relies on conformational changes in the selectivity filter (Abderemane-Ali et al., 2019). Given the proximity of the tryptophan residues in the structure of the selectivity filter to the Ca^2+^ coordinating residues, we considered whether these mutations also alter CDI. At baseline, Ca_V_1.3 channels exhibit strong CDI (Fig. 5A, rose curve) quantified as the fraction of excess inactivation with Ca^2+^ versus Ba^2+^ as charge carrier measured following 300ms of depolarization (*CDI* = 1 – *r*_300,Ca_ / *r*_300,Ba_). Here, CDI measurements were performed at ambient CaM levels. Analysis of DI W[+2]A, DII W[+2]A, and DIII W[+2]A mutants revealed minimal changes in CDI (Fig. 5A-C). As DIV W[+2]A mutation enhances VDI, comparison of inactivation kinetics with Ca^2+^ versus Ba^2+^ as permeant ion is insufficient to quantify CDI. To obviate this limitation, we estimated CDI as excess inactivation of Ca^2+^ currents with endogenous CaM compared to with Ca^2+^-binding deficient CaM_1234_ (see Methods). The latter maneuver disables CDI and provides an estimate of average VDI with Ca^2+^ as permeant ion (Fig. 5D, blue). Thus measured, the DIV W[+2]A mutant showed a modest but statistically-significant reduction in CDI (*p* < 0.001, Fig. 5E).

**Figure 5.**
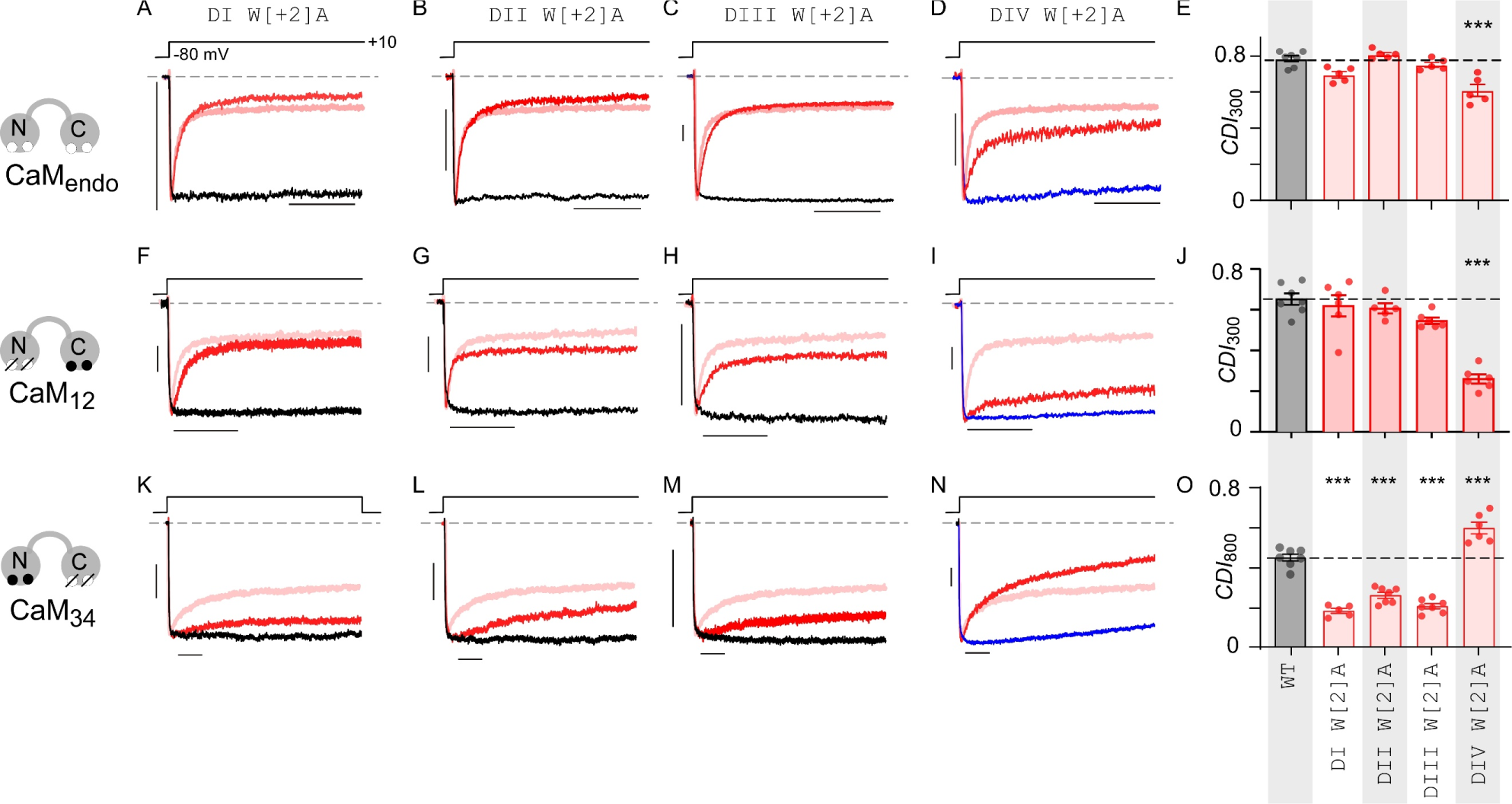
Structural biparitioning of CDI mediated by distinct CaM lobes. **(A)** Exemplar currents show strong CDI of DI W[+2]A mutation as evident from accelerated current decay observed with Ca^2+^ (red) versus Ba^2+^ (black) currents evoked in response to a 300 ms test pulse. Ca^2+^ current decay of wild-type Ca_V_1.3 is shown as rose trace to facilitate comparison. Here, measurement obtained in the presence of endogenous CaM and therefore net CDI includes contributions of both N- and C-lobes of CaM. **(B – C)** DII W[+2]A and DIII W[+2]A mutations minimally perturb CDI. Format as in panel A. **(D)** Ca_V_1.3 DIV W[+2]A mutation partially diminishes CDI. Here CDI is measured by comparing inactivation in Ca^2+^ with endogenous CaM versus CaM_1234_ co-expressed (blue trace) **(E)** Population data confirms a mild reduction in CDI observed with W[+2]A mutation. p<0.001 by Tukey’s multiple comparisons test. Each bar and error, mean ± s.e.m. *n* = 7 cells for Ca_V_1.3 wild-type and *n* = 5 for all mutant channels. (**F-J**) CDI mediated by C-lobe of CaM is selectively diminished by Ca_V_1.3 DIV W[+2]A mutation. Here, the component of C-lobe CDI is isolated by co-expression of CaM_12_. For DIV W[+2]A mutant, we compare inactivation with CaM_12_ compared to CaM_1234_. Red trace, Ca^2+^ current; Black trace, Ba^2+^ current; Rose trace, Ca^2+^ current from Ca_V_1.3 WT co-expressed with CaM_12_. Each bar and error, mean ± s.e.m. *n* = 6 cells (Ca_V_1.3 wild-type), *n* = 5 cells (DI W[+2]A), *n* = 5 cells (DII W[+2]A), *n* = 6 cells (DIII W[+2]A), and *n* = 7 cells (DIV W[+2]A). ***p<0.001 by Tukey’s multiple comparisons test. (K-O) CDI mediated by N-lobe of CaM is diminished by Ca_V_1.3 DI W[+2]A, DII W[+2]A, and DIII W[+2]A mutations. However, DIV W[+2]A increases N-lobe CDI. Here, N-lobe CDI is isolated by co-expression of CaM_34_. Red trace, Ca^2+^ current; Black trace, Ba^2+^ current; Rose trace, Ca^2+^ current from Ca_V_1.3 WT co-expressed with CaM_34_. For DIV W[+2]A mutant, we compare inactivation with CaM_34_ versus when CaM_1234_ is co-expressed (blue trace). Each bare and error, mean ± s.e.m. *n* = 5 cells (Ca_V_1.3 wild-type), *n* = 7 cells (DI W[+2]A), *n* = 7 cells (DII W[+2]A), *n* = 6 cells (DIII W[+2]A), and *n* = 7 cells (DIV W[+2]A). ***p<0.001 by Tukey’s multiple comparisons test.

As Ca_V_1.3 CDI is mediated by both N- and C-lobes of CaM (Yang et al., 2006), we next considered whether pore domain mutations may differentially impact regulation by the distinct CaM lobes. To evaluate this possibility, we measured C-lobe CDI by co-expressing mutant CaM_12_ with Ca^2+^ binding to its N-lobe disabled, or N-lobe CDI in the presence of CaM_34_ whose Ca^2+^ binding to the C-lobe is disabled. Indeed, we found that DI W[+2]A, DII W[+2]A, and DIII W[+2]A mutants had minimal or no effect on C-lobe CDI (Fig. 5F-H, 5J). However the DIV W[+2]A mutation strongly inhibited C-lobe CDI (Fig. 5I-J). By contrast, we observed that N-lobe CDI was sharply diminished for DI W[+2]A, DII W[+2]A, and DIII W[+2]A mutants (Fig. 5K-M, 5O), while the DIV W[+2]A mutant exhibited an increase in CDI (Fig. 5N-O). These findings highlight the asymmetric contribution of pore domain residues in differentially coupling CDI triggered by distinct CaM lobes.

### DIV W[+2] mutation enhances inactivation of TS-linked Ca_V_1.2 variant

Human mutations in Ca_V_1.2 cause a severe multisystem disorder known as Timothy Syndrome (TS), marked by an increased likelihood for cardiac arrhythmia and neurological and neurodevelopmental deficits. Mechanistically, these channelopathic variants are well established to exhibit reduced inactivation (both VDI and CDI) resulting in prolonged Ca^2+^ entry that delays action potential repolarization and disrupts Ca^2+^ homeostasis (Barrett and Tsien, 2008; Dick et al., 2016; Calorio et al., 2019). As such, identifying mechanisms that reverse this change would be beneficial from the perspective of devising new therapeutics. Since DIV W[+2]A mutation in Ca_V_1.3 enhanced VDI, we probed whether the analogous mutation in a TS Ca_V_1.2 variant could reverse deficits in VDI observed for mutant channels. We considered the G406R variant, known to exhibit a strong functional phenotype. Consistent with previous studies (Barrett and Tsien, 2008; Dick et al., 2016), we found that this variant exhibited reduced both VDI and CDI compared to wild-type Ca_V_1.2 channels (Fig. 6A-B). Introduction of the DIV W[+2]A mutation to the G406 variant resulted in a marked enhancement in VDI (Fig. 6C), restoring overall inactivation to near wild-type levels. This finding suggests that enhancing the conformational flexibility of the SF may be a potential approach to reverse the pathophysiological reduction in inactivation linked to TS.

**Figure 6.**
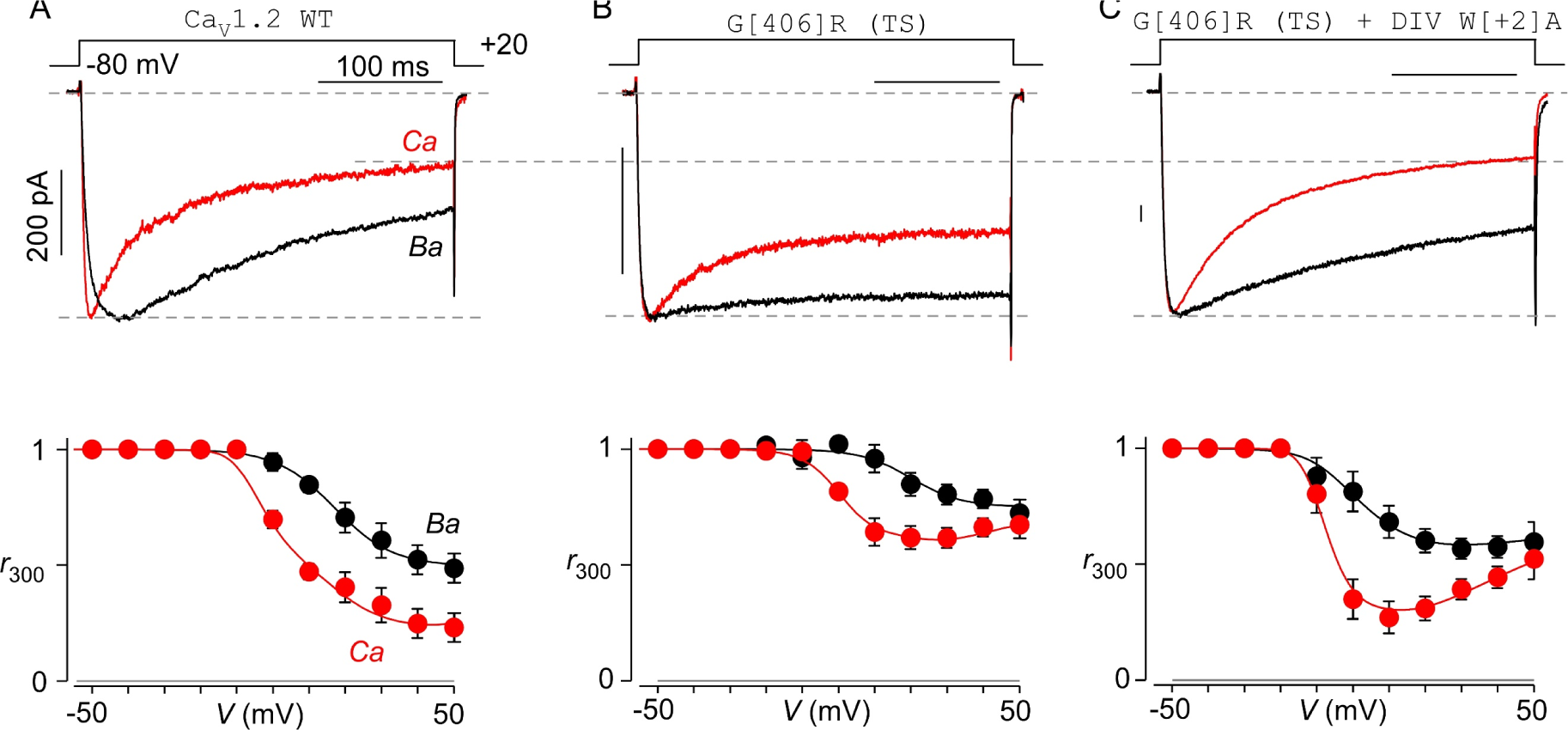
SF domain mutation enhances inactivation of Timothy Syndrome associated Ca_V_1.2 mutant channel. **(A)** Top, wild-ty pe human Ca_V_1.2 channel exhibits robust CDI and VDI. Black trace, Ba^2+^ current; red trace, Ca^2+^ current. Bottom, population data show baseline inactivation of Ca_V_1.2 measured as *r*_300_, the fraction of peak current remaining following 300 ms, with either Ca^2+^ (red) or Ba^2+^ (black) as charge carrier. Each symbol, mean ± s.e.m. *n* = 5 cells. (**B**) Ca_V_1.2 G406R mutant linked to Timothy syndrome shows reduced VDI and CDI. Format as in panel A. *n* = 6 cells. (**C**) Introducing the DIV W[+2]A mutation on TS-linked CaV1.2 G406R background enhances VDI. This finding suggests that increasing the conformational flexibility of DIV SF may be a potential approach to reverse TS pathophysiology. *n* = 6 cells.

## Discussion

Ca_V_ channel inactivation is a physiologically consequential and mechanistically rich ion channel feedback regulation that serves to limit excess Ca^2+^ entry into neurons and cardiomyocytes (Minor and Findeisen, 2010; Ben-Johny and Yue, 2014). Here, we probed the functional importance of the SF in tuning both VDI and CDI. Mutagenesis of an ultra-highly conserved W[+2] residue (Payandeh and Minor, 2015) in domains I through IV revealed an unexpected asymmetric role of the pore domain in tuning channel inactivation. First, we found that mutating the W[+2] residue in DIV but not DI through DIII enhanced VDI. MD simulations showed that DIV W[+2]A mutation increased the local conformational flexibility of the pore domain, hinting at a distinct role for the SF gate in tuning VDI. Although the W[+2] residue is symmetric in all four domains, the asymmetric effect of mutating DIV W[+2] residue on VDI stems from distinct differences in neighboring residues that stabilize the interdomain interfaces. Second, in depth analysis of CDI revealed a structural bifurcation of CaM signaling, whereby the DIV mutation selectively diminished CDI mediated by the C-lobe of CaM, while mutations in DI – DIII preferentially reduced N-lobe CDI. Taken together, these findings demonstrate that the pseudo-tetrameric arrangement of the eukaryotic Ca_V_ channel pore enables asymmetric conformational changes that are fundamentally important for its distinct inactivation mechanisms.

For VDI, a hinged-lid mechanism has been proposed based on multiple lines of evidence (Stotz et al., 2000; Dafi et al., 2004; Tadross et al., 2010): (1) Chimeric analyses of Ca_V_1/2 channels pointed to the involvement of both the DI S6 segment and the intracellular DI-DII linker in tuning inactivation kinetics (Stotz and Zamponi, 2001; Stotz et al., 2004). (2) Kinetics of VDI is dependent on the specific auxiliary Ca_V_β subunit associated with the channel DI-DII linker, with membrane tethered Ca_V_β subunits slowing VDI (Olcese et al., 1994; Chien et al., 1996). (3) Mutagenesis of the presumed S6 transmembrane receptor site also alters VDI (Stotz and Zamponi, 2001; Dafi et al., 2004; Tadross et al., 2010). (4) Exogenous expression of DI-DII linker as a peptide accelerates VDI (Cens et al., 1999). Structurally, the DI-DII linker forms a continuous helix with the S6 gate and disruption of the helix by a poly-glycine linker promotes VDI, findings that are inconsistent with a hinged-lid model (Findeisen and Minor, 2009; Wu et al., 2016). Recent studies instead point to the involvement of the SF for VDI of Ca_V_ channels, with select mutations in the SF diminishing VDI (Abderemane-Ali et al., 2019). In this study, the slow baseline VDI of Ca_V_1.3 enabled us to discern the potential role of the SF using a gain-of-function approach by introducing mutations to upregulate VDI. Our results suggest that the SF interface between DIII and DIV may be a previously unrecognized locus for VDI. Although the W[+2] residue is symmetrically present in all four domains, only the DIV W[+2]A mutation increased local conformational flexibility and enhanced VDI. Further mutagenesis of this residue showed that the kinetics of VDI varied depending on the bulkiness of the residue at the DIV [+2] position, with bulkier residues slowing inactivation kinetics. This asymmetric effect of the DIV mutation reflects distinct differences in residues at the DIII-DIV interdomain interface (Fig. 7A). Specifically, the W[+2] residue interacts with a T[-2] residue, that is present in all four domains. Beyond this, the DI-DII, DII-DII, and DIV-DI interdomain interfaces are also stabilized by additional interactions between a either Q[-5] or R[-5] and a Y[+7] residue from the neighboring domain. However, this additional stabilizing interaction is absent in the DIII-DIV interface, with a T[-5] in the DIII and an L[+7] in the DIV locations. Consequently, the DIV W[+2]A mutation destabilizes the SF conformation. Consistent with this possibility, the DIV W[+2]A / DIII T[-5]Q double mutation partially reversed the increase in VDI. Taken together, the inter-domain interaction of DIV W[+2] with neighboring residues may be envisioned as a brake that stabilizes the SF conformation allowing sustained ion influx during prolonged depolarization.

**Figure 7.**
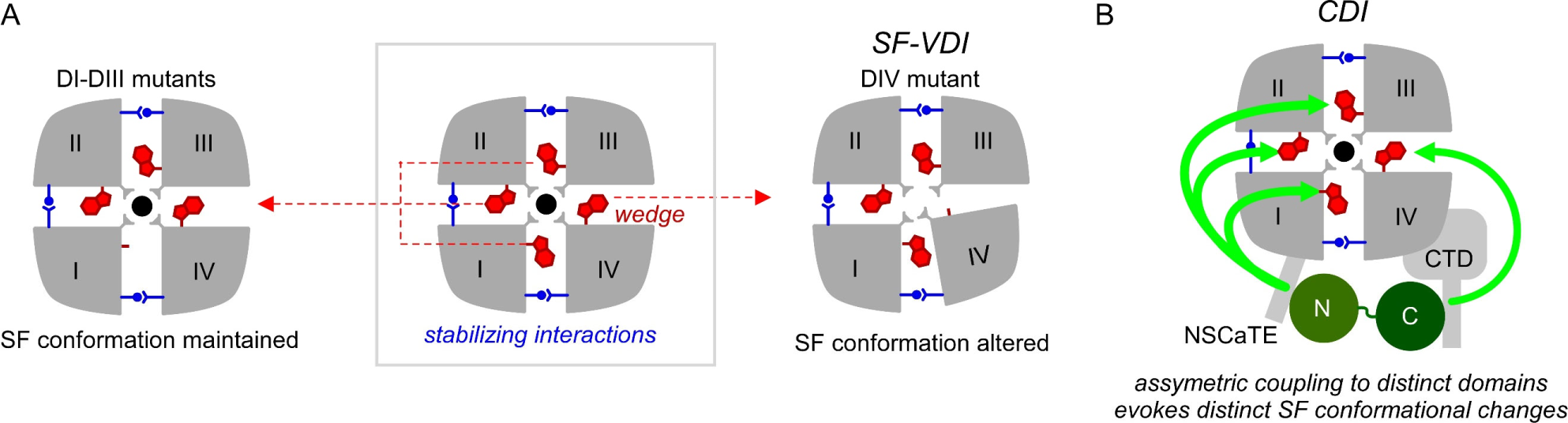
Asymmetric effect of SF mutations on Ca_V_1.3 VDI and CDI. (A) Schematic shows asymmetric effect of the SF W[+2]A mutations in evoking VDI of Ca_V_1.3. The interdomain interface between DIII and DIV is stabilized primarily by the conserved W[+2] residue which may serve as a wedge. Disruption of this interaction destabilizes the local conformation. By contrast, the interdomain interface between DI-DII, DII-DII, and DI-DIV are stabilized by both the W[+2] residue and through additional inter-domain contacts. Consequently, SF conformation is maintained even with the W[+2]A mutation in DI through DIII. (B) For CDI, the W[+2] residue likely plays a role in transducing SF conformational rearrangements. The N- versus C-lobe of CaM appear to be asymmetrically coupled to distinct SF domains, likely evoking distinct SF conformational changes.

For Ca_V_1.3, mutating residues in the distal S6 domain below the bundle crossing has also been shown to enhance VDI (Tadross et al., 2010). However, this form of VDI exhibited a conventional Boltzmann-shaped voltage-dependence and its magnitude was susceptible to further modulation by Ca_V_β subunits. By contrast, VDI uncovered by SF mutations exhibited an atypical U-shaped voltage-dependence and was insensitive to modulation by the Ca_V_β subunits. The disparate functional signatures suggest that the two forms of VDI are distinct. Interestingly, Ca_V_2.2 channels have been shown to also undergo preferential intermediate closed-state inactivation with a similar U-shaped voltage-dependence (Patil et al., 1998). Recent cryo-EM studies identified a cytosolic helical segment knowns as W-helix unique to Ca_V_2 DII-DIII linker that occludes the ion conduction pathway (Dong et al., 2021; Gao et al., 2021). Either deletion or disruption of this domain by alternative splicing was shown to diminish preferential intermediate closed state inactivation (Dong et al., 2021). It is possible that altered conformational changes in the SF of Ca_V_2.2 may also contribute to this process.

CDI is initiated by CaM-dependent conformational changes in the cytosolic domain of Ca_V_ channels (Budde et al., 2002; Halling et al., 2006; Minor and Findeisen, 2010; Ben Johny and Yue, 2015). Ca^2+^ binding to the individual N- and C-lobes of CaM can trigger distinct forms of channel regulation (Lee et al., 2000; DeMaria et al., 2001; Liang et al., 2003), a phenomenon known ‘functional bipartition’ of CaM (Kink et al., 1990; Saimi and Kung, 1994). For Ca_V_1.3, the N- and C-lobe of CaM evokes kinetically distinct forms of inactivation (Yang et al., 2006; Dick et al., 2008). Molecularly, these effects are initiated by a Ca^2+^-dependent switching of CaM interaction with distinct channel interfaces – the Ca^2+^-bound N-lobe engages the *NSCaTE* domain on the channel amino-terminus (Dick et al., 2008; Simms et al., 2013), while the C-lobe associates with the carboxy-terminus and evokes a conformational change (Kim et al., 2004; Black et al., 2005; Van Petegem et al., 2005; Asmara et al., 2010; Ben-Johny et al., 2013; Banerjee et al., 2018). How these disparate changes in the cytosolic domains are coupled to the pore domain are not fully understood. Two end-stage mechanisms of CDI have been proposed: (1) an allosteric change in the activation gate corresponding to a change in modal gating (Tadross et al., 2010) or (2) an altered selectivity filter conformation (Abderemane-Ali et al., 2019). First, single channel recordings show that Ca^2+^/CaM regulation switches channels from a high activity gating with ‘flickery’ openings to a low activity gating mode with ‘sparse’ openings (Imredy and Yue, 1994). Furthermore, both engineered and human disease-linked mutations in the S6 gate that enhance channel activation proportionately weaken CDI, all consistent with an allosteric mechanism. (Tadross et al., 2010). Second, Ca_V_1.2 inactivation is sensitive to extracellular Ca^2+^ and CDI alters propensity for pore blockade by Gd^3+^, suggesting involvement of the SF (Babich et al., 2005; Babich et al., 2007). More concretely, mutations in the SF alter CDI. Specifically, SF DII D[+1]A diminishes CDI of various Ca_V_1/2 channels without any apparent effect on VDI (Abderemane-Ali et al., 2019). Our present findings further corroborate an essential role for the SF in mediating CDI and uncover an unexpected asymmetry in this process. The DIV W[+2]A mutation that destabilizes the SF, evokes contrasting effects on CDI mediated by distinct CaM lobes (Fig. 7B). While this mutation enhances N-lobe CDI, it also suppresses C-lobe CDI. By contrast, mutating corresponding W[+2] residues in DI-DIII revealed a selective effect on N-lobe CDI but not C-lobe CDI. This divergent effect suggests that the specific inter-domain interactions of the W[+2] residue in the SF may be involved in transduction of cytosolic conformational changes to the end-stage mechanism. Of note, MD simulations did not show a marked increase in conformational flexibility with DI-DIII W[+2]A mutations. However, these simulations did not include the channel carboxy- and amino-terminal domains which are essential for CaM regulation. It is possible that CaM rearrangement in the cytosolic domains may evoke additional changes in the SF domain. These findings also suggest that there may be distinct SF conformational changes associated with N- versus C-lobe CDI. How does one reconcile an allosteric mechanism evident from single-channel studies with a change in SF conformation? One possibility emerges from studies of Kcsa channels where altered SF conformation has been shown to result in shifts in modal gating (Chakrapani et al., 2011). The activation gate of these channels is also coupled to the SF gate (Cuello et al., 2010; Heer et al., 2017), suggesting that the two mechanisms may overlap. Similar findings have also been reported with other ion channels (Boiteux et al., 2014; Kopec et al., 2019; Coonen et al., 2020).

From a pathophysiological perspective, channelopathic mutations in both Ca_V_1.2 and Ca_V_1.3 are known to reduce both VDI and CDI (Dick et al., 2016; Ortner et al., 2020; Bamgboye et al., 2022a). This deficit in inactivation is thought to result in sustain Ca^2+^ influx that is proarrhythmic. From a pharmacological perspective, traditional small molecule blockers of Ca_V_1 channels have been largely inadequate as a therapy. First, mutant channels often exhibit a reduced affinity for many small molecule blockers (Bamgboye et al., 2022b). Second, small molecule Ca_V_1 inhibitors impact the peak Ca^2+^ current, which in turn inhibits the overall Ca^2+^ influx causing adverse negative inotropic effects (Angelini et al., 2021). Instead, Ca_V_1 gating modifiers that enhance inactivation have been proposed as a new and potentially more effective class of antiarrhythmic drugs. In this study, we found that introduction of the DIV W[+2]A mutation can in fact enhance the VDI of TS-linked mutant channels to near wild-type levels. This finding suggests that either destabilizing or increasing the conformational flexibility of DIV SF domain may suffice to upregulate Ca_V_1 inactivation and potentially reverse pathophysiology of Timothy Syndrome.

Of broader relevance, the W[+2] residue is highly conserved across eukaryotic Ca_V_, Na_V_, and NALCN leak channels and prokaryotic BacNa_V_, suggesting that asymmetric SF conformational changes may constitute an ancient and general mechanism for channel inactivation (Payandeh and Minor, 2015). For Na_V_1.4, the DI W[+2]C mutation was shown to inhibit slow inactivation while mutations of analogous residues in DII-DIV had minimal effect (Balser et al., 1996). In like manner, mutating the DIV A[0] residue that forms part of the SF of Na_V_1.4 introduces an ultra-slow component of inactivation with a U-shaped voltage-dependence, much like the Ca_V_1.3 DIV W[+2]A mutation (Todt et al., 1999). For the homo-tetrameric bacterial Na_V_ channels, asymmetric pore collapse is thought to be the underlying structural mechanism for inactivation (Pavlov et al., 2005; Payandeh et al., 2012). The W[+2] residue serves as an important anchor for the SF that allows hydrogen bonding interactions with T[-2] residues of neighboring subunits (Payandeh and Minor, 2015). This interaction has been proposed to be important to allow SF conformational changes to be influenced by neighboring subunits (Payandeh and Minor, 2015). Structurally, slow-inactivation of BacNa_V_ channels increases the conformational flexibility of the selectivity filter, resulting in a reorientation of the pore helices that change the inter-subunit cavity volumes and accessibility of lipids (Chatterjee et al., 2018).

Overall, this study furnishes new insights into the role of the SF in VDI and CDI of Ca_V_1.3 and it uncovers how asymmetric changes in the SF allow structural bifurcation of CaM signaling, a scheme that may be broadly relevant (Saimi and Kung, 2002; Liang et al., 2003; Ben-Johny et al., 2014).

## Acknowledgements

We thank Dr. Ryan Mahling, Dr. Nourdine Chakouri, Dr. Ivy Dick, Dr. Po Wei Kang, and Dr. Filip van Petegem for helpful feedback. We thank Audrey Kochiss for technical support. This study is supported by funding from NINDS (R01 NS110672) to MBJ, American Heart Association Predoctoral Fellowship (award ID 835091) to P.J.D, a grant from the Austrian Science Fund (FWF) P35618 to B.E.F, and Austrian Academy of sciences APART-MINT postdoctoral fellowship to M.L.F.Q. and the Austrian Science fund (P34514). The content is solely the responsibility of the authors and does not necessarily represent the official views of the National Institutes of Health or the American Heart Association.

## Author Contributions

P.J.D., M.L.F-Q., K.L., B.F., and M.B-J. designed research, P.J.D., M.L.F-Q., A.L.K. and M.B-J. performed research, acquired and analyzed data, P.J.D., M.L.F-Q., K.L., B.F., and M.B-J., contributed new reagents / analytic tools; M.B-J., funding acquisition; P.J.D., M.L.F-Q., and M.B-J made figures and wrote the original draft; and all authors revised the manuscript. The computational results presented here have been achieved in part using the Vienna Scientific Cluster (VSC). We thank PRACE for awarding us access to Piz Daint at CSCS, Switzerland.

## Materials and Methods

### Molecular Biology

Mutagenesis Ca_V_1.3 was performed using truncated variant of rat Ca_V_1.3 (AF370009.1), Ca_V_1.3_Δ1626_ as previous published (Tadross et al., 2010; Ben-Johny et al., 2013). Briefly, we first PCR amplified subsegments of wild-type Ca_V_1.3 containing SF domains of domain I and domain II (R1, flanked by restriction enzyme sites BsiWI and Eco47III), domain III (R2, flanked by restriction enzyme sites Eco47III and BglII), and domain IV (R3, flanked by restriction enzyme sites BglII and XbaI) respectively. These subsegments were inserted into the pCRBlunt II-TOPO vector using the Zero Blunt TOPO PCR cloning kit (Invitrogen). Point mutations in respective Ca_V_1.3 subsegments were then generated using Quikchange II kit (Agilent). We generate W357A and W718A, we used R1 plasmid, while R2 plasmid was used for W1114A and R3 plasmid for W1404A, W1404F, W1404V, and W1404T mutations. Subsequently, the mutated segments were ligated into the wild-type Ca_V_1.3 following restriction digest with the specific enzymes for a given subsegment. Ligates were then transformed into either XL10-Gold Ultracompetent Cells (Agilent) or DH5α Competent Cells (Thermo Fisher), plated and cultured in selective LB broth. DNA was extracted and purified from cultures using either QIAprep Spin Miniprep Kit (Qiagen) or GeneJET PCR Purification Kit (Thermo Fisher). Sanger sequencing was used to verify each mutant. For T1107Q/W1404A double-mutant, we first generated the T1107Q mutation using Quikchange II (Agilent) in the R2 subsegment. Subsequently, this region was ligated into W1404A mutant channel following restriction digest using Eco47III and BglII.

### Cell Culture and Transfection

For whole-cell electrophysiology, HEK293 cells were cultured on glass coverslips in 60-mm dishes and transfected using a calcium phosphate method. We applied 2–8 μg of cDNA encoding the desired channel α_1_ subunit (WT or engineered variant), along with 4 μg of rat β subunit (β_1b_, β_2a_, β_3_ or β_4_) and 4 μg of rat brain α_2_δ (NM012919.2). In experiments that required CaM expression, we transfected 2-4 μg of cDNA encoding CaM WT or mutant variants (CaM_12_, CaM_34_, or CaM_1234_). To enhance expression, cDNA for simian virus 40 T antigen (0.5 μg) was co-transfected. Electrophysiology recordings were done at room temperature 1–3 days after transfection.

### Whole-cell Recordings

Whole-cell voltage-clamp recordings for HEK293 were collected at room temperature using an Axopatch 200B amplifier (Axon Instruments). Glass pipettes (World Precision Instruments, MTW 150-F4) were pulled with a horizontal puller (P97; Sutter Instruments Co.) and fire-polished (Microforge, Narishige, Tokyo, Japan), resulting in 1–3-megaOhm resistances, before series resistance compensation of 70%. Internal solutions contained 135 mM CsMeSO_3_, 5mM CsCl_2_, 1mM MgCl_2_, 4mM MgATP, 10 mM HEPES, 10 mM BAPTA, adjusted to 290-295 mOsm with CsMeSO_3_ and pH 7.4 with CsOH. The external solutions contained 140 mM TEA-MeSO_3_, 10 mM HEPES (pH 7.4), either 40 mM CaCl_2_ or BaCl_2_. This external solution composition was chosen based on previous studies to ensure that local Ca^2+^ signals are saturating to drive maximal local CDI (Tadross et al., 2008). For experiments using Na^+^ as charge carrier, we used external solution containing 140 mM NaCl, 10 mM HEPES (pH 7.4), 10 mM TEA-MeSO_3_ and 0.5 mM EGTA. Solutions were adjusted to 300 mosM with TEA-MeSO_3_ and pH 7.4 with TEA-OH. For CDI and VDI measurements, we used a family of test pulses to −50 to +50 mV with repetition intervals of 20-120 s, from a holding potential of −80 mV. Custom MATLAB (Mathworks) software was used to determine peak current and fraction of peak current remaining after either 300 ms (r300) of depolarization or 800 ms (r800) of depolarization. Residual currents after depolarization were measured at +10 mV after an initial pulse of +10 mV for 15 ms followed by a family of test pulses from −60 to +50 mV for 800 ms. Custom MATLAB (Mathworks) software was used to determine peak current and I_2_/I_1_ ratio. As W[1404]A mutation itself enhanced VDI, quantifying CDI (i.e. the CaM driven component of inactivation) by comparing inactivation with Ba^2+^ versus Ca^2+^ are charge carrier is imprecise. Accordingly, we estimated baseline VDI for W[1404]A mutation in the presence of Ca^2+^ by coexpressing CaM_1234_ (*r*_300,CaM1234_ = *I*_300ms_ / *I*_peak_ and *r*_800,CaM1234_ = *I*_800ms_ / *I*_peak_, where *I*_peak_ is the peak current, and *I*_300_ and *I*_800_ are current levels following 300 ms or 800 ms of depolarization respectively. We then measured total inactivation (CDI and VDI) with Ca^2+^ as charge carrier with endogenous CaM, CaM_12_ or CaM_34_ to isolate total, C- and N-lobe components of CDI, respectively (*r*_300,Ca_ = *I*_300ms_ / *I*_peak_ and *r*_800,Ca_ = *I*_800ms_ / *I*_peak_ with Ca^2+^ as charge carrier). Subsequently, CDI was estimated as CDI_300_ = 1-*r*_300,Ca_/ *r*_300,CaM1234_) or CDI_800_ = 1-*r*_800,Ca_/ *r*_800,CaM1234_). For all other mutations, we observed minimal VDI. As such CDI was measured as CDI_300_ = 1-*r*_300,Ca_/ *r*_300,Ba_ and CDI_800_ = 1-*r*_800,Ca_/ *r*_800,Ba_.

### Molecular Dynamics Simulations

As starting structures for our simulations, we used the recently published Cryo-EM structure of the Cav1.3 Ca^2+^ channel complex (PDB accession code: 7UHG) (Yao et al., 2022). The structures of the W1404A and the W718A mutants were derived from the wild-type model by replacing the mutated residue followed by a local energy minimization using MOE (Molecular Operating Environment, version 2020.09, Molecular Computing Group Inc., Montreal, Canada). The C-terminal and N-terminal parts of each domain were capped with acetylamide (ACE) and N-methylamide to avoid perturbations by free charged functional groups. The structure model was embedded in a plasma membrane consisting of POPC (1-palmitoyl-2-oleoyl-sn-glycero-3-phosphocholine) and cholesterol in a 3:1 ratio, using the CHARMM-GUI Membrane Builder (Jo et al., 2008; Jo et al., 2009). Water molecules and 0.15 M CaCl_2_ were included in the simulation box. Energy minimizations of wild-type and mutant structures in the membrane environment were performed. The topology was generated with the LEaP tool of the AmberTools21, using force fields for proteins and lipids, ff14SBonlysc and Lipid14, respectively (Dickson et al., 2014). The wild-type and mutant structures were gradually heated from 0 to 300 K in two steps, keeping the lipids fixed, and then equilibrated over 1 ns. Then molecular dynamics simulations were performed for 3x300 ns, with time steps of 2 fs, at 300 K and in anisotropic pressure scaling conditions. Van der Waals and short-range electrostatic interactions were cut off at 10 Å, whereas long-range electrostatics were calculated by the Particle Mesh Ewald (PME) method. PyMOL was used to visualize the key interactions and point out differences in the wild-type and mutant structures (The PyMOL Molecular Graphics System, Version 2.0 Schrödinger, LLC).

## Supplementary Information

**Figure S1.**
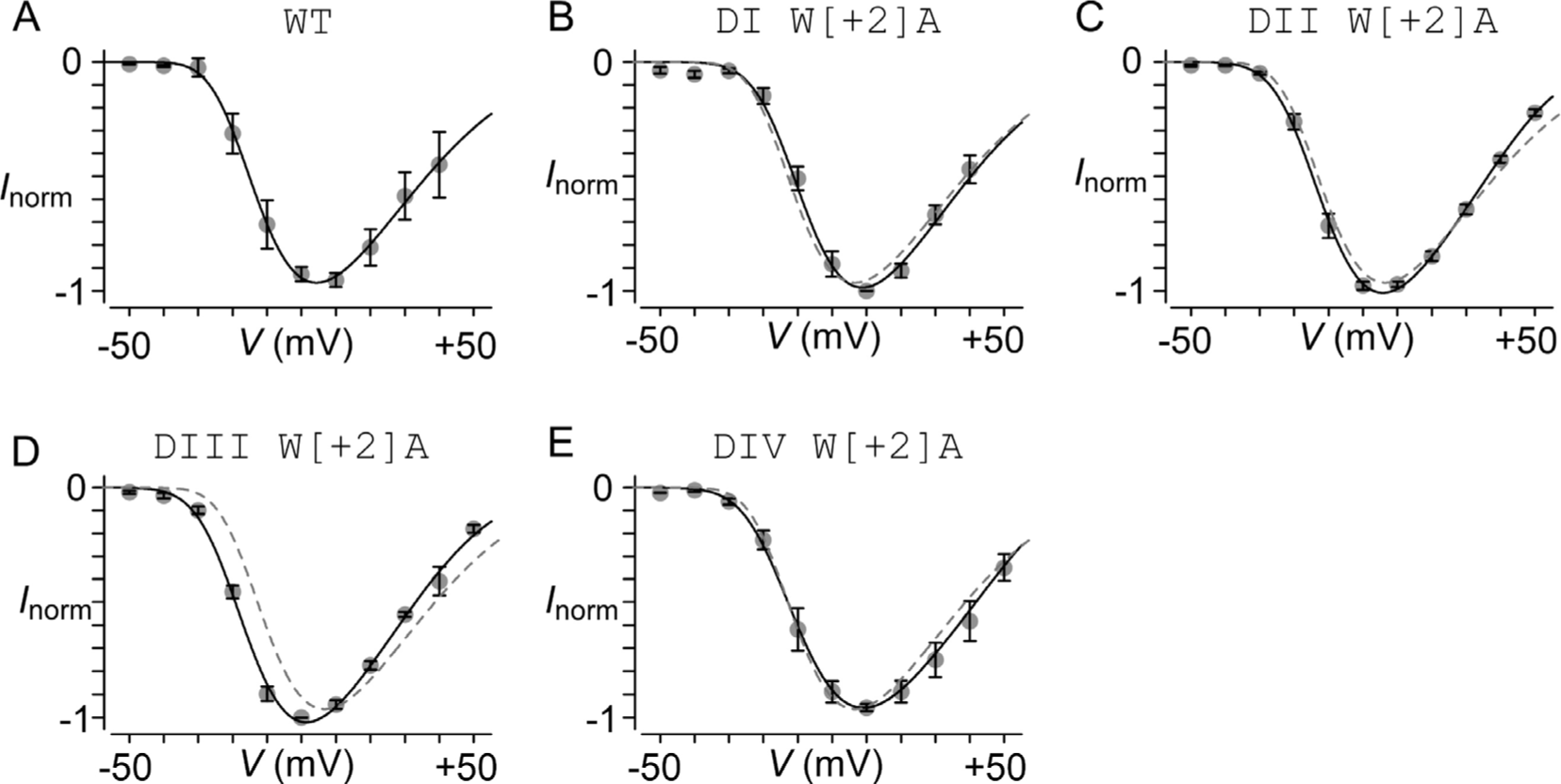
Ca_V_1.3 DI-DIV SF mutations minimally perturb channel activation. (**A**) Normalized current voltage-relationship for WT Ca_V_1.3. Each dot, mean ± s.e.m. *n* = 5 cells. (**B – E**) Normalized IV relations for DI/W[+2]A (panel B, *n* = 5 cells), DII/W[+2]A (panel C, *n* = 6 cells), DIII/W[+2]A (panel D, *n* = 5 cells), and DIV/W[+2]A (panel E, *n* = 7 cells) mutants. In all cases, the change in channel half-activation potential (*V*_1/2_) was less than 5 mV.

**Figure S2.**
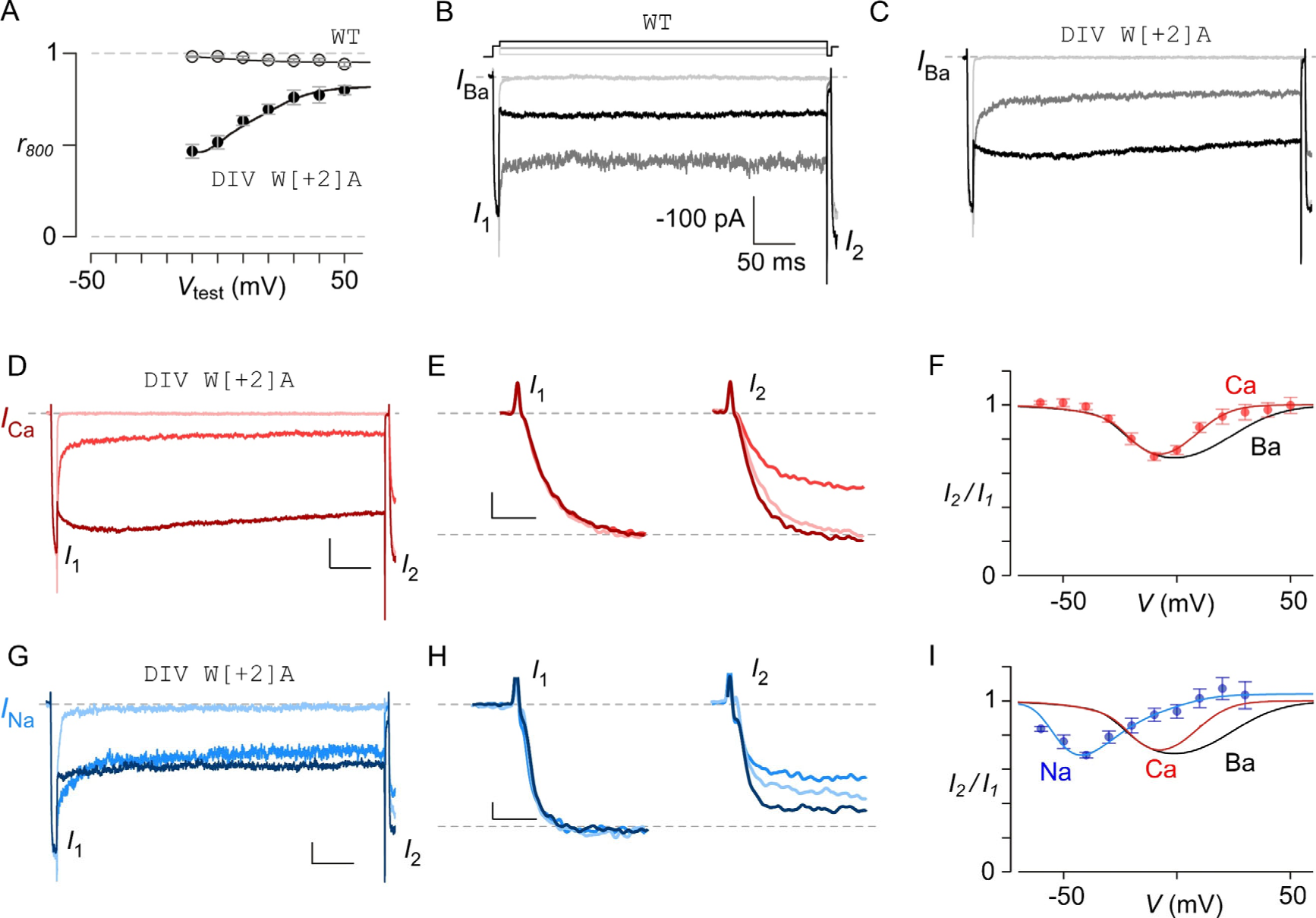
Extended Biophysical characterization of DIV W[+2]A mutation. (**A**) Population r800 data plotted as a function of test-pulse potential reveals reduced inactivation at higher voltages. Each dot, mean ± s.e.m. *n* = 7 cells. (**B**) A two-pulse protocol is used to dissect voltage-dependence of inactivation. A brief 15 ms prepulse to 10 mV quantifies available current prior to inactivation. Subsequently, a family of 800 ms voltage steps (test pulse) is used to evoke steady-state inactivation and a 15 ms postpulse is then used to quantify the current remaining following VDI. The extent of inactivation is quantified as the ratio of peak currents during post-pulse to pre-pulse. Exemplar currents at three different test pulse potentials (light gray, −50 mV; gray, 0 mV; and black +50 mV) are shown. Exemplar trace for wild-type Ca_V_1.3. Further analysis in main text Fig. 2C, E. (**C**) Exemplar trace for DIV W[+2]A mutation. Further analysis in main text Fig. 2D, F. (**D – F**) VDI of DIV / W[+2]A probed in the presence of Ca^2+^ as charge carrier. Panel D, exemplar trace. Panel E, comparison of Ca^2+^ currents during pre-pulse (left, *I*_1_) versus post-pulse (right, *I*_2_). Note that the current magnitude during the post-pulse is maximally reduced at intermediate voltages. Panel F, population data shows U-shaped dependence of inactivation with Ca^2+^ as charge carrier (red). Black trace, relationship with Ba^2+^ as charge carrier reproduced from Fig. 2F. Each dot, mean ± s.e.m, *n* = 8 cells. (**G-I**) VDI of DIV / W[+2]A mutation with Na^+^ (but no divalents) as charge carrier. Format as in Panel D-F. *n* = 7 cells.

